# A neural representation of invisibility: when stimulus-specific neural activity negatively correlates with conscious experience

**DOI:** 10.1101/2020.04.20.051334

**Authors:** Matthew J Davidson, Will Mithen, Hinze Hogendoorn, Jeroen J.A. van Boxtel, Naotsugu Tsuchiya

**Author notes:** Equal contribution (Jeroen J.A. van Boxtel, Naotsugu Tsuchiya). Author contacts,.

## Abstract

Although visual awareness of an object typically increases neural responses, we identify a neural response that increases prior to perceptual *disappearances*, and that scales with the amount of invisibility reported during perceptual filling-in. These findings challenge long-held assumptions regarding the neural correlates of consciousness and entrained visually evoked potentials, by showing that the strength of stimulus-specific neural activity can encode the conscious absence of a stimulus.

**Significance Statement:** The focus of attention and the contents of consciousness frequently overlap. Yet what happens if this common correlation is broken? To test this, we asked human participants to attend and report on the invisibility of four visual objects which seemed to disappear, yet actually remained on screen. We found that neural activity increased, rather than decreased, when targets became invisible. This coincided with measures of attention that also increased when stimuli disappeared. Together, our data support recent suggestions that attention and conscious perception are distinct and separable. In our experiment, neural measures more strongly follow attention.

## Introduction

Research on the neural basis of conscious perception has almost exclusively focused on an awareness of a stimulus that is present. This research has a long history in showing that becoming aware of a stimulus leads to increases in neural responses (e.g., ^1–3^). These findings go together with an often implicit assumption that the disappearance of a stimulus, and awareness of its absence, should lead to weaker neural responses. Here we show using electroencephalography (EEG), that the disappearance of a stimulus can positively correlate with increases in neural activity. This, in turn, casts doubt on the direct relationship that has been observed between neural activity and conscious perception, at least when measured with current neuroimaging tools such as EEG. We argue that it is not conscious perception that has traditionally been correlated with these neural measures, but attention, and that neural activity and conscious perception will only positively correlate when increases in attention lead to increased visibility.

To break the common correlation between attention and visibility we employed a perceptual filling-in (PFI) paradigm, in which the probability of a stimulus disappearing (i.e. being filled-in) increases, rather than decreases, with attention ^4,5^. For example, attending to shared colours ^4^, or forms ^5^ among targets increases their reported rates of disappearance. During PFI, the disappearing stimuli are perceptually replaced by the surrounding visual background, despite their continued physical presence. The conscious awareness of target-presence is therefore negatively correlated with attention in this paradigm. Existing theories propose that a combination of activity in retinotopic visual cortex ^6,7^, and higher-level association areas ^5,8,9^ initiate and maintain these periods of invisibility ^10^.

To obtain large-scale entrained neural measures of stimulus disappearance and reappearance we employed EEG in a novel multi-target multi-response PFI paradigm ^11^. EEG measures consisted of steady-state visually evoked potentials (SSVEPs), which are periodic cortical responses that entrain to rhythmic visual stimuli. SSVEPs have a high signal-to-noise ratio (SNR), and as a result have been widely adopted in the basic visual, cognitive, and clinical neurosciences ^12,13^. Importantly, they have been extensively used to measure the focus of attention without overt responses ^14,15^, as well as the contents of consciousness ^1–3^.

To capture neural markers of attention and/or consciousness during PFI, we isolated the specific neural activity contributed by both targets and visual surrounds while human participants continuously reported on the visibility of multiple targets. We achieved this by dynamically updating our visual display within a target boundary at 15 Hz (F1), and the target surrounds at a rate of 20 Hz (F2; **Movie 1**, https://osf.io/kdrz3/**, Figure 1**). This approach entrained neural populations sensitive to the target or surrounds at different frequencies, and allowed us to directly monitor changes to the time-course of target or surround-specific neural activity. These unique neural signals are expected to anti-correlate if the SSVEP signal is related to visibility, because target invisibility is inextricably linked to surround visibility, due to filling-in. However, these signals are expected to correlate positively when they are related to attention, because both the disappearance of the target, and filling-in of the surround, are expected to attract attention.

**Figure 1.**
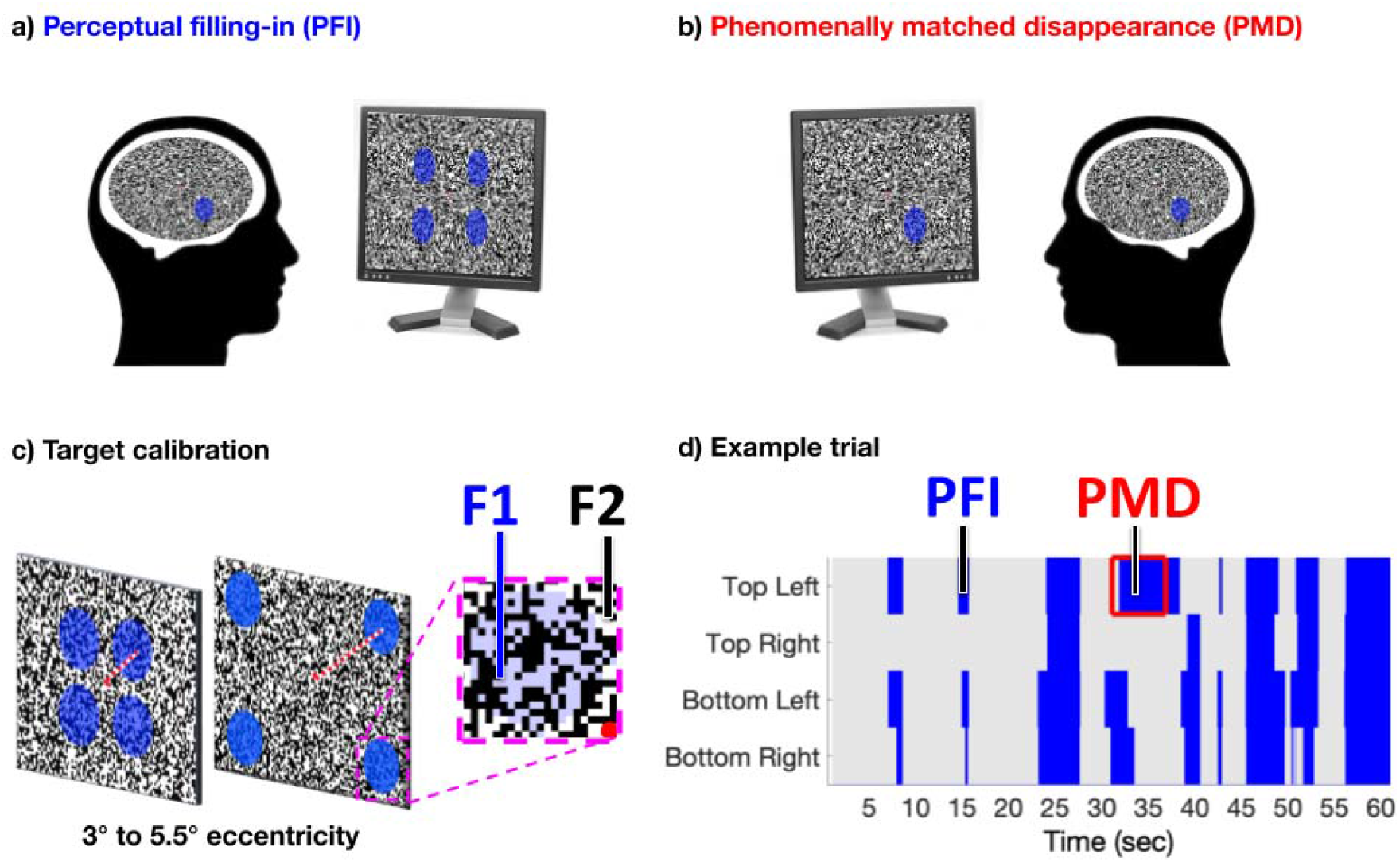
Task Design. a) Four peripheral targets, defined by their blue-colour, were superimposed over a dynamic texture to induce perceptual filling-in (PFI). b) Phenomenally-matched disappearances (PMD) were embedded within each trial, to mimic the subjective quality of PFI, during which one to four targets were physically removed from screen. c) The eccentricity of targets was calibrated per participant, using a custom brain-computer-interface, real-time SSVEP display (see Methods). This procedure optimised PFI, target (F1) and surround (F2) frequency-tagged responses. d) Each of 48 trials lasted 60 s, during which participants reported on the visibility of all four peripheral targets, using four unique buttons.

## Results

### A neural marker of subjective invisibility

Our participants (*N*=16) reported on the subjective invisibility of four simultaneously presented peripheral targets, located in separate visual quadrants (**Figure 1**). We analyzed the time-course of target (F1) and surround (F2) SNR during PFI, aligned to button-press at target disappearance, and button-release at reappearance. Remarkably, we found that the strength of target-specific neural responses *increased* prior to the subjective report of a target becoming *invisible*. Compared to target reappearance, this effect was significant from −0.74 s before, through to 1.53 s after subjective report (*p*_cluster_ <.001), indicating that target-SNR is negatively correlated with the contents of conscious perception during PFI. We next sorted all instances of PFI by the amount of simultaneous button press, measuring the magnitude of subjective invisibility (i.e., amount of PFI; **see Methods**). We found that increases in the amount of PFI (i.e., decreased visibility) significantly increased target-SNR (linear-mixed effects, likelihood ratio test: χ^2^(1) = 11.60, *p* = 6.58 x 10^−4^; **Figure 2c-blue**), confirming that target-SNR increased as a measure of target invisibility during PFI. Unlike target-SNR, surround-SNR tracked the contents of phenomenology: it increased during the filling-in of targets, compared to their reappearance, from −0.99 to 1.28 s around report (*p*_cluster_ < .001; **Figure 2e**), and it linearly increased with the amount of PFI (χ^2^(1) = 19.63, *p* = 9.41 x 10^−6^; **Figure 2f-black**). Thus, target and surround signals are positively correlated during PFI, suggesting they are not related to visibility. For a discussion of the timing differences between target- and surround-SNR, see **Supplementary Results**.

**Figure 2.**
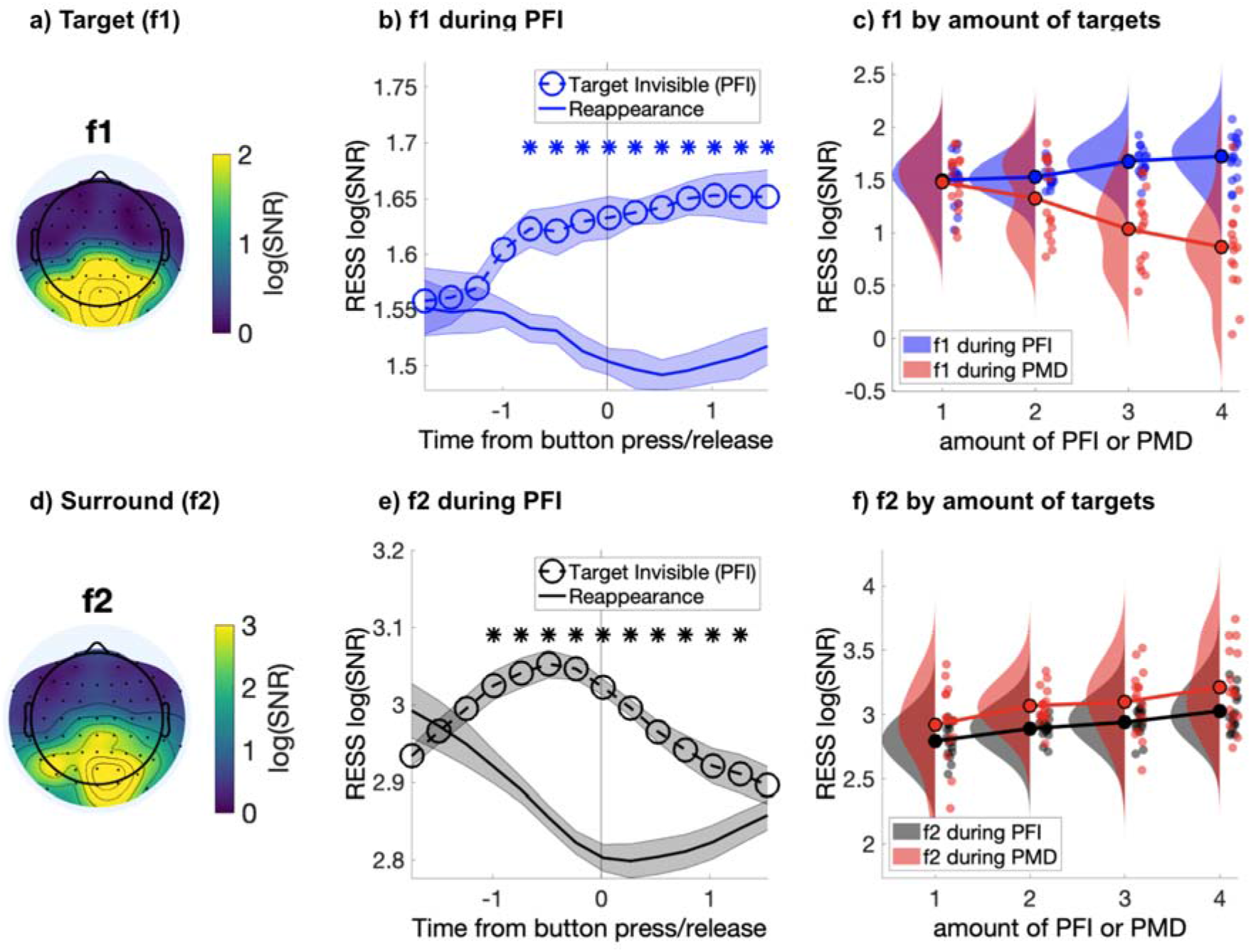
Target and surround-specific neural responses during PFI. a) Topoplot of whole-trial target-SNR at f1 (15 Hz). b) Target-SNR negatively correlates with visibility of the target (i.e., increases with invisibility and decreases with visibility). c) The negative correlation for target-SNR is unique to PFI, during physically matched disappearance (PMD), target-SNR positively correlates with the amount of visual targets. Shading in b and c corresponds to 1 SEM corrected for within-subject comparisons (Cousineau, 2005). d) Topoplot of surround-SNR at f2 (20 Hz). e) Surround-SNR increases during PFI, matching phenomenology as targets become interpolated by their surroundings. f) Increased experience of the surround during both PFI and PMD positively correlate with surround-SNR.

### Increased SNR during invisibility is unique to targets during PFI

We also analysed the time-course of SNR during Phenomenally Matched Disappearance (PMD) periods (**Figure 1b**). These were events embedded within every trial, during which the phenomenal disappearance experienced during PFI was mimicked by physically removing targets from the screen (**see Methods**). During PMD, target-SNR significantly decreased (χ^2^(1) = 18.35, *p* = 1.84 x 10^−5^; **Figure 2c-red**), rather than increased as during PFI. No dissociation between PMD and PFI periods was observed for surround-SNR, which significantly increased with increasing amounts of PMD (χ^2^(1) = 7.37, *p* = .007; **Figure 2f-red**), as it did for PFI, confirming that only during PFI, when periods of invisibility were endogenously generated, did specifically target-sensitive neural populations increase in SNR.

### Visibility negatively correlates with a neural measure of attention

An increased SSVEP response to invisible targets is incompatible with the long-standing interpretation that SSVEP amplitude indexes stimulus visibility, or indeed the contents of consciousness (e.g. ^16^). Instead, our data suggest that SSVEP responses are primarily due to other factors, such as fluctuations in the allocation of attention (e.g.^14,17^). Indeed, attention increases the likelihood of PFI in psychophysical experiments^4,5^, and we also observed a synergistic effect between multiple PFI events (**Supplementary Results, Supplementary Figure 5**), which may indicate enhanced perceptual grouping due to attention ^11,18^.

To test whether the SSVEP changes in our data were indeed due to attention, we performed an exploratory analysis, in which we used evoked responses in the alpha band (8-12 Hz, baseline corrected from −2.5 to −1.5 s prior to button press) as an index of attention during PFI (**Figure 3a**). Evoked alpha responses are frequently used as a measure of attention ^19–23^ Event-related reductions in alpha are commonly interpreted as a measure of enhanced cortical excitability under top-down control ^24,25^, and correlate with the subjective intensity of attentional effort ^26^. In our paradigm, we hypothesised an event-related reduction in alpha would be consistent with a role of attention during PFI.

**Figure 3.**
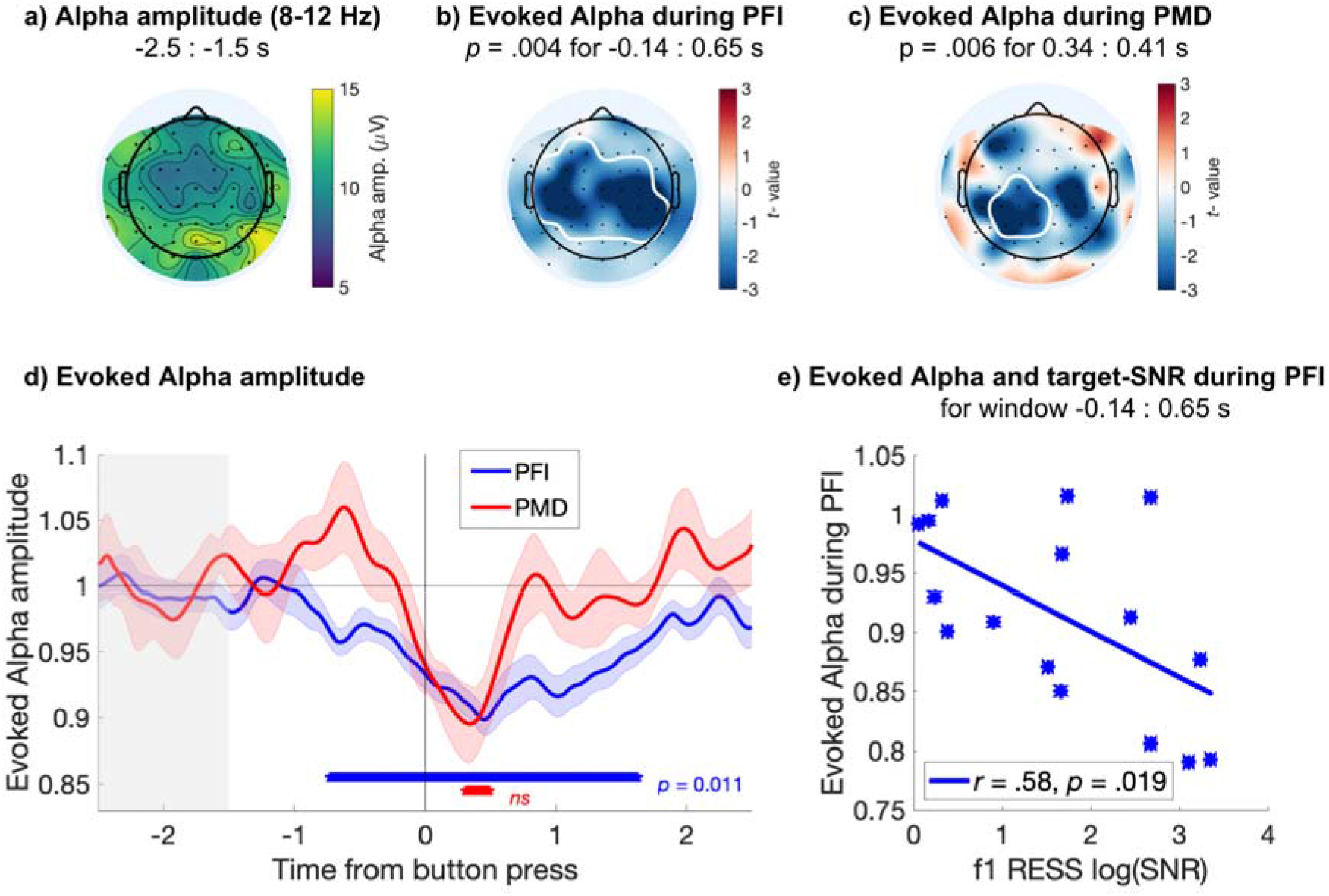
Evoked responses in the alpha band. Centro-parietal alpha amplitude decreases during PFI. a) Spatial topography of alpha band (8-12 Hz) amplitude during baseline (−2.5 to −1.5 s before button press). b) PFI- and c) PMD-evoked reductions in alpha amplitude, with significant clusters outlined in white (electrode-time cluster corrected). d) Time course of evoked alpha during PFI (blue) and PMD (red), averaged over the significant electrodes marked in b and c, respectively. Shading shows 1 SEM, corrected for within-participant comparisons (Cousineau, 2005). Blue asterisks mark the significant time-window for the evoked alpha during PFI, averaged over significant electrodes in b (temporal cluster corrected), p = .011 at FDR q = .05. (The effect during PMD did not survive FDR corrections for multiple comparisons here). e) Each participant’s alpha amplitude averaged over the significant time-window shown in b (−0.14 to 0.65 s), plotted against the target SNR at f1 during the same time-window.

With a cluster-based permutation test (**see Methods**), we confirmed a significant difference between baseline alpha amplitude (averaged −2.5 to - 1.5 prior to button press), and alpha amplitude evoked during PFI (**Figure 3b**; *p_cluster_* = .004, electrode-cluster-time cluster corrected; ^27^). This condition difference corresponded to a centro-parietal cluster of electrodes, and reduction in alpha during PFI from - 0.14 s through to 0.65 s after button press. During PMD, a smaller centro-parietal cluster of electrodes, and smaller, later latency range (0.35 to 0.41 s) was significant. (**Figure 3c**; *p_cluster_* = .006, 0.35 to 0.41 s; electrode-cluster-time cluster corrected; ^27^). Cluster-based permutation tests should be interpreted with caution, and used to infer only about the overall statistical contrast, and not to infer the location or latency of a significant effect ^27,28^. To grasp a clearer picture of the significant effect we identified, we also display the time-course of the evoked reductions in alpha during PFI and PMD, averaged over the electrode clusters depicted in **Figure 3b and 3c**, respectively. When comparing the evoked alpha during PFI and PMD to the prestimulus baseline, alpha amplitude significantly reduced from approximately −0.77 to 1.58 s during PFI (**Figure 3d**; *p* < .011 at FDR q = .05). During PMD, no significant reduction in alpha was retained after false-discovery rate corrections (FDR q = .05). Across participants, mean evoked alpha also negatively correlated with increases to target-SNR during PFI (*r* = −0.58, *p* = .019, **Figure 3e**), in further support of attention driving PFI dynamics in our paradigm. No correlation was found between evoked alpha reductions and target-SNR during PMD, (*r* = −0.34, *p* = .222).

## Discussion

By demonstrating that visibility negatively correlates with object-specific neural responses, while positively correlating with our neural measure of attention (reduced alpha-amplitude), our results challenge contemporary theories of conscious perception. For example, first-order theories of conscious perception ^29,30^ propose that the strength of activity within content-specific visual processing areas determines the vividness of that experience in conscious vision. Our results challenge this account, as stimulus-specific neural activity positively correlated with subjective *invisibility*. Our findings are also difficult to accommodate within the global neuronal workspace theory (GNWT) of consciousness ^31^. According to GNWT, attention typically acts as a gateway to conscious access, enhancing stimulus representations for wider broadcasting, and awareness positively correlates with the strength of sensory-specific information^32^. In our data, the focus of attention did not correspond with the vividness of an object in consciousness, and neural measures of attention increased with the invisibility of objects. As more targets disappeared and presumably drew attention, both the duration of their absence and strength of target-SNR increased. To explain our findings, some aspects of GNWT may need modification.

Alternatively, theories of consciousness that consider interactions among neurons or neural populations and their states to be related more directly to phenomenology are more compatible with our paradigm. Currently, one such candidate is integrated information theory ^33–35^, where the informational structure of neural states can be computed as neural correlates of perception ^36^, without prior assumptions regarding stimulus strength or attention.

Overall, our results reinforce that studying the neural basis of consciousness is fraught with difficulty, especially since the contents of consciousness and focus of attention so frequently overlap ^37,38^. This overlap helps to explain why the SSVEP has been interpreted as a measure of the content of conscious perception for over 50 years, since Lansing first correlated the contents of a subjective report with the amplitude of rhythmic brain responses ^2^. In this study, we leveraged a perceptual paradigm in which attention decreases conscious visibility during PFI, thereby dissociating the two processes. We have measured a neural signature of this dissociation, which may lead to an explanation of more puzzling phenomena, such as attentionally enhanced stimulus disappearance ^4,5^, while strongly suggesting that the neural correlates of attention and perception are dissociable.

We show increased neural responses to the disappearance of stimuli, which challenges long-held beliefs about the correlation between neural measures and consciousness, by suggesting that the brain may actively represent disappearing, or absent stimuli. Previous research has suggested neurons in the prefrontal cortex encode abstract decisions for the absence of stimuli ^39^, but our report is the first to show entrained neural responses arising from the visual cortex which represent the invisibility of disappearing stimuli in human participants. This suggests that the awareness of an absence is important to the brain, and perhaps that further breakthroughs will follow an acknowledgement of how absence awareness is a valid conscious experience, which may support the neural mechanisms and correlates of consciousness.

## Methods & Materials

### Participants

19 healthy adults (12 female, 19-40 years, M = 26.95, SD = 7.63) with normal or corrected-to-normal vision participated in this study. They were recruited via convenience sampling from students at Monash University, provided written informed consent prior to taking part, and were paid 20 AUD per hour of their time (approximately 3 hours total). Participants with self-reported sensitivity to flickering stimuli were excluded. Ethics approval was obtained from the Monash University Human Research Ethics Committee (MUHREC #CF12/2542 - 2012001375).

### Apparatus and stimuli

Participants sat approximately 50 cm from a computer monitor (size 29 x 51 cm, resolution 1080 x 1920 pixels, subtending 32 x 54° visual angle, refresh rate 60 Hz). The background surrounding our target stimuli (**Figure 1**) was refreshed at a rate of 20 Hz by randomly selecting one of 100, pre-calculated, random patterns generated at the start of the experiment. The random patterns were made up by dividing the screen into squares of 10 x 10 pixels (0.3 x 0.3° visual angle), with each square’s luminance randomly set to either black or white. Visually, this procedure appeared similar to random noise, with an equal proportion of black and white texture. At the centre was a 5 mm (0.57° visual angle) diameter red dot serving as a fixation point.

One circular target was presented in each of the four visual quadrants (top left, top right, bottom left, and bottom right). Targets subtended 6.08° visual angle, within which the white background squares were instead coloured as blue/purple (RGB value of 205, 205, 255). There were no line-contours at target boundaries, as target boundaries were defined by the pixelated squares of the background they were overlaid on (**Figure 1c**). As a result, during PFI, target regions became perceptually indistinguishable from the surrounding background texture. The refresh rate of texture within the target regions was set to 15 Hz, with target eccentricity individually calibrated per participant to evoke optimal PFI and SSVEP strength (see **Figure 1c**, and Pre-experiment SSVEP calibration of target eccentricity).

### EEG acquisition

Throughout each session, EEG was recorded with 64 active electrodes arranged according to the international 10-10 system (BrainProducts, ActiCap). Electrode impedances were kept below 10 kΩ prior to experimentation, and recorded using the default reference (FCz) and ground electrode (AFz) via Brainvision recorder software (sampling rate =1000 Hz, offline bandpass of 0.5-70 Hz). All EEG data was stored for offline analysis using custom Matlab scripts (Ver: R2016b), as well as the EEGLab ^41^, Chronux ^42^, and Fieldtrip ^43^ toolboxes. Prior to experimentation, the BCILAB toolbox ^44^ was also implemented to calibrate real-time SSVEP strength (see below).

### Pre-experiment SSVEP calibration of target eccentricity

Finding optimal conditions for SSVEP and PFI paradigms is fraught with difficulty, as the features which enhance each are in direct opposition: in order to optimize SSVEPs, large and foveally-positioned targets are preferred ^13^, while to optimize PFI, small and peripherally-positioned targets are required ^6,45^. We solved this problem using a custom brain-computer-interface (BCI) procedure.

The purpose of the calibration procedure was to find a target eccentricity at which PFI readily occurred, while evoking observable peaks in the EEG spectra at the target flicker frequency (15 Hz). Targets were initially centred close to fixation (3° visual angle from centre). Participants were instructed to fixate a central dot, and were asked to report on PFI. If participants reported perceptual filling-in, they were then asked to describe their experience more explicitly. Depending on their subjective reports, we adjusted the eccentricity of the targets along the diagonal (from 3 - 5.5°, in steps of 0.3°) so that their perceptual experience was described as “appearing and disappearing often, for a few seconds at a time”. All participants reported that perceptual filling-in occurred. For six participants, optimal perceptual filling in was not achieved, describing their experience as “relatively invisible, only occasionally reappearing.” These participants were retained for all analyses.

Concurrently with these manipulations, the power and log(SNR) spectra at POz were displayed in real time. Due to the computational demands of presenting frequency-domain EEG spectra in real time, no inferential statistics were used to define adequate SSVEP strength. Readily observable 15 and 20 Hz peaks in the EEG spectra were taken as evidence of frequency-tagging at face value. In the absence of readily observable peaks in the real time EEG spectra, target eccentricity was reduced, and the process was repeated. If tagging was unsuccessful at all settings (*n*=2), the largest and most central target position at which any perceptual filling-in occurred was adopted. This was under the assumption that repetition over many trials may still result in a frequency-tag, which was confirmed in our analyses. **Supplementary Figure 1** displays the final calibrated target eccentricity across participants (*N*=19).

### Experimental procedure

Participants were instructed with the following script: “Fixate on the red dot. If you perceive that any of the four targets has completely disappeared, press the button corresponding to that target and hold it down for as long as you perceive that target to be absent. If more than one target vanishes simultaneously, try to report on them all as accurately as possible.” Specifically, they were instructed to press keys ‘A’, ‘Z’, ‘K’, and ‘M’ on a traditional QWERTY keyboard, assigning them to the upper left, bottom left, upper right, and bottom right targets, respectively. After calibration, each participant completed one 60-second practice trial, followed by 48 self-paced, experimental trials which were also one minute long. Participants took mandatory breaks of 3-5 minutes every 12 trials, while EEG recordings were paused and channel impedances were monitored.

### Phenomenally matched disappearance (PMD) periods

Each trial included one randomly generated phenomenally-matched disappearance (PMD) period of between 3.5 and 5 s (drawn from a uniform distribution), during which one to four targets were physically removed from the screen. Participants were not informed of these physical disappearance periods before the experiment. To mimic the frequency of genuine PFI, PMD did not occur in the first 10 s of each trial ^46^. The order of all PMD periods were randomized for each participant, as were the location of removed targets in the case of one, two and three targets. These physical disappearance periods enabled us to monitor how well participants were able to report on the visibility of four simultaneously presented targets, and additionally served as a control condition for comparison with the neural signals evoked by genuine PFI.

### Participant and trial exclusion based on PMD periods

We have previously demonstrated that the use of PMD periods can identify participants who are unable to report on four simultaneous targets, as well as experimental trials which are unsuited for further analyses ^11^. In particular, whether a participant accurately reported the PMD period on time (via button press) was used to estimate that participant’s attention to the task.

Unlike our previous study ^11^, we tailored our stimulus so that PFI happens very often across all participants. Thus, it was often the case that the physical removal of a target (PMD onset) occurred for targets which have already disappeared from consciousness due to genuine PFI. Such occurrences were more frequent for those participants experiencing greater amounts of PFI. To obtain accurate estimates of how accurately and quickly participants responded to the physical removal of a target we estimated the baseline button-press likelihood per individual participant, by performing a bootstrapping analysis (with replacement). To perform this bootstrap, for any PMD onset in trial T at time S (seconds), we randomly selected a trial T’ (T=T’ was allowed) and we epoched [S-2, S+4] button press data at corresponding PMD target locations in T’.

We repeated this for all available trials (T=1…48) to obtain a single bootstrapped set of 48 trials per participant. This procedure was repeated 1000 times, and the mean button press time-course for each 48-trial set was retained as the null-distribution of button-press likelihood, over time, per participant. **Supplementary Figure 2d (grey)** shows the bootstrapped likelihood of button press for a single participant. We then used the null distribution z-scores of ±1.96 as the 95%CI for each participant (after first converting the data using the logit transformation due to a violation of normality). We defined the reaction time to PMD-onset as the first time point after which each participant’s median button-press data exceeded the top CI of their null-distribution likelihood. We excluded participants (*n*=3) with no PMD-onset reaction time in the first 2 s (i.e., [0, S+2]). We note that for two of these participants, it appeared that buttons were consistently released at PMD-onset - potentially indicating buttons were released during PFI rather than pressed as per instructions. For the remaining participants, the mean reaction time to respond to PMD onsets, and thus the disappearance of a peripheral target was 0.68 s (*SD* = 0.31).

Having identified which participants could successfully indicate target disappearance based on their button press data (*N*=16), we continued to remove any trials in which a PMD was not correctly detected from subsequent analysis. This ensured that throughout each trial, participants were accurately reporting on PFI, and that all retained data was indicative of complete attention to target regions. To identify individual trials for exclusion, we regarded a PMD period as failed if the corresponding button was not pressed for at least 50% of the allowed response time window. This window was the duration of the PMD (3.5 to 5 s). **Supplementary Figure 2c** shows a histogram of the number of rejected trials per participant (overall, only 16 trials were rejected) across 16 retained participants.

### Simultaneous PFI and location-shuffling analysis

To examine whether an increasing number of filled in targets (nPFI; *n* = 0, 1, 2, 3, 4) affected PFI characteristics, we performed a shuffling analysis to create a null distribution that removed the temporal correlation between targets (Davidson et al., 2020). This analysis tests whether simultaneous PFI occurs more often that can be expected by chance. Specifically, we created 1000 shuffled trials per participant, that could contain the button-press data for each location from any trial throughout their experimental session (e.g. one shuffled trial could consist of top left button press time course from trial 10, top right from trial 43, bottom left from trial 12, bottom right from trial 2). We selected a random trial for each location, allowing multiple locations from one trial to be included to form one shuffled trial (i.e., bootstrap with repetition). This shuffling procedure conserves the total amount of PFI recorded, while ensuring that the button-press data at a given location could come from any independent trial. If target disappearances during PFI were independent, then destroying the temporal correlation in shuffled data should not matter, and shuffled and experimental data would look similar. We repeated our behavioural analysis (as detailed above) on this shuffled data, with the results displayed in **Supplementary Figure 5**.

We also performed quadratic model-fit analyses to compare the magnitude and direction of the fitted coefficients for observed and shuffled data. For this analysis, we retained the coefficient (β; 2nd order polynomial) from the fit to our observed data as our observed statistic. We also fit the same quadratic model to each of the shuffled data sets (*n* = 1000) and used these coefficient values as a null distribution to compare with the observed β. If the observed β exceeded the top 95% of the null distribution, we regarded the quadratic fit for the observed data as significant at *p* < 0.05.

### EEG preprocessing

After data collection, noisy channels were identified using a modified version of the PREP pipeline ^47^. We omitted the bad-by-RANSAC criterion that identifies correlated channel groups which deviated from other channels. This was necessary as frequency-tagging elicits responses in localised, often highly correlated channel clusters. Bad channels were then spherically interpolated (channels rejected per participant *M* = 5.70, *SD* = 0.62). After channel rejection and interpolation, whole-trial EEG data were re-referenced to the average of all electrodes, and linearly detrended before being downsampled to 250 Hz. We then applied a Laplacian transform to improve the spatial selectivity of our data. **Supplementary Figure 3** shows the whole trial spectrum from electrode POz, as well as topographical distribution of frequency-tagged components.

### SSVEP analysis via rhythmic entrainment source separation (RESS)

SSVEP topography can vary based on individual participant anatomy, the entrained neural network ^48^ as well as based on the frequency of flicker selected (as shown in the above figure). As such, we applied a spatiotemporal filter called rhythmic entrainment source separation (RESS), to reduce the distributed topographical response at SSVEP frequencies to a single component time-series ^49^. The RESS single component is a weighted average from across all channels, which can be analyzed in the time-frequency domain instead of relying upon the selection of a single or multiple channels based on post-hoc data inspection. Specifically, RESS works by creating linear spatial filters tailored to maximally differentiate the covariance between a signal flicker frequency and neighbourhood frequencies, thereby increasing the signal-to-noise ratio at the flicker frequency. We constructed RESS spatial filters from 64-channel EEG, by applying a narrow-band Gaussian (full-width at half maximum = 0.5 Hz) filter to the original EEG data in the frequency-domain, centered at the target, surround and IM frequency, respectively (and independently). As the frequency-neighbourhood across different signals would contain different amounts of simultaneous flicker, we proceeded by selecting broadband neural activity to construct reference covariance matrices. Comparing signal to broadband activity has previously been shown to allow the reconstruction of SSVEP signals using RESS ^11,49^.

RESS spatial filters were constructed per participant per frequency (target flicker and harmonics; 15, 30, 45 Hz, surround flicker and harmonics 20, 40, 60 Hz; Intermodulation components; 5, 25, 35 Hz), using epoched data from all time-windows −3000 to −100 ms and 100 ms to 3000 ms around button press/release, avoiding PMD periods. We avoided the 200 ms around button press/release to optimize RESS filters in the absence of motor-evoked activity. Each filter was fitted without distinguishing whether targets were disappearing or reappearing due to button press or release, in order to reduce the possibility of overfitting these condition comparisons. After application of the RESS spatial filters, we reconstructed the time course of SSVEP log(SNR) from the RESS component time-series as described below (SSVEP SNR calculation). With RESS, we were able to focus our analysis on a single component time-series per frequency of interest, without arbitrarily selecting a single channel or averaging channels, eliminating the need for corrections for multiple comparisons across channels.

### SSVEP Signal-to-Noise Ratio (SNR) calculation

In the SSVEP paradigm, we compute the signal-to-noise ratio (SNR) at each frequency, which in logarithmic scale corresponds to log of the power at each frequency minus the mean log power across the neighbourhood frequencies. Throughout this paper, the neighbourhood used in SNR calculation always excludes the frequency half-bandwidth nearby f Hz, which is calculated with a formula: f = (k+1) /2T, where k = the number of tapers used in time-frequency decomposition and T= temporal window (seconds) of the data.

For our moving-window SNR analyses, we first computed the single-taper (k=1) spectrogram using a 2.5 s window, with a half-bandwidth of 0.4 Hz, and shifted the window in a step size of 250 ms. We then compared the signal at f to neighbourhood [f-2.5, f-0.5] and [f+0.5, f+2.5]. For our whole-trial SNR analysis with a half-bandwidth of 0.017 Hz, we compared the signal at f to neighbourhood [f-0.12, f-0.06] and [f+0.06, f+0.12].

### Evoked responses in the alpha band

We used a two-step analysis process to calculate the relative alpha amplitude during all PFI and PMD periods. First, after all PFI and PMD periods were epoched and preprocessed as described above, we bandpass filtered the time series with a narrow-band Gaussian filter (8-12 Hz, full-width at half maximum = 2 Hz). Second, to the bandpass filtered signals, we applied the Hilbert transform to compute the alpha-band amplitude as the envelope of the Hilbert-transformed filtered signal. Third, to obtain evoked responses in the alpha band, within participants, we divided the time course of the amplitude by the average over the period −2.5 to −1.5 s prior to button press (grey window shown in **Figure 3d**) for each participant.

To identify electrode-cluster-time points, in which the difference between baseline and evoked alpha responses were significant, we implemented non-parametric cluster-based permutation tests (Fieldtrip ^27,43^; ft_timelockstatistics.m function). Specifically, using all electrodes, we set a minimum spatial cluster size of three neighbours, and a threshold for cluster identification at *p* = .05 (uncorrected). Spatiotemporal clusters of test-statistics (*t*-scores from dependent samples *t*-tests) which exceeded *p* < .05 (uncorrected) were then identified. We then corrected for multiple-comparisons using Monte Carlo permutation tests (1000 repetitions). In each permutation, we exchanged the data label between conditions at random within participants, and repeated the above procedure to identify spatiotemporal clusters on these permuted data, and obtained a null distribution for clustered-test statistics (*t*-scores). By comparing the null distribution to the observed cluster statistic, we finally obtain the cluster-level p-value corrected for multiple-comparisons ^27^.

### Event-by-Event image analysis of button press and log(SNR)

We performed image-based event-by-event analysis ^50^, to investigate whether the changes in log(SNR) may reflect the amount of PFI. This image-based analysis is necessary due to variations in the frequency and duration of reported PFI per participant.

To accommodate these differences, we first sorted all PFI events in descending order, based on the sum of buttons pressed over a 3 s period. We used the period [0, +3] relative to button press for PFI disappearances, and [-3, 0] relative to button release for PFI reappearances. This integral of the number of buttons pressed we term “the amount of PFI” ^11^. After sorting based on the amount of PFI, we then resampled participant data along the trial dimension (y-axis) to normalize trial counts to 100 trials for each participant. This process of resampling along the trial dimension was repeated for the event-by-event time course of log(SNR), except the order of trials was predetermined by the corresponding button-press per participant. A schematic pipeline for this entire procedure is displayed in **Supplementary Figure 4**. Finally, to quantify whether changes in log(SNR) occur with an increasing amount of PFI, we grouped trials when the amount of PFI was between 0 and 1, 1 and 2, 2 and 3 or greater than 3. The results have been visualized using the raincloud plots statistical package ^51^.

### Statistical analysis – EEG

To assess the significance of SSVEP peaks in the EEG spectra, we corrected for multiple comparisons with a False Discovery Rate (FDR) q = .05 (Benjamini et al, 2006) (Supplementary Figure 3).

We corrected for multiple comparisons in SNR time-series (Figure 2 b and e) using non-parametric (temporal) cluster-based corrections ^27^. Specifically, we first detected any temporally contiguous cluster by defining a significant time point as p < .05 uncorrected. Then, we concatenated the contiguous temporal timepoints with p < .05 and obtained a summed cluster-level test statistic for the cluster. The sum of observed test-statistics (e.g., t scores) in a temporally contiguous cluster were then retained for comparison with a permutation-based null distribution. To create the null distribution, we repeated the procedure of searching for and retaining contiguous time-points which satisfied the p < .05 (uncorrected) cluster criterion, after first shuffling the condition labels 2000 times. For within-participant comparisons, this amounts to randomly permuting the averages for each condition within each participant. From each of the 2000 repetitions, we obtained the maximum sum of cluster-level test statistics, which served as a null distribution. We regarded the effect to be significant if the original summed cluster-level statistics exceeded the bottom 95% of the null distribution of the summed statistics (as p_cluster_ < .05).

## Supplementary Results

### A spatial interaction between target locations increases PFI duration when targets disappear together

We investigated behavioural responses during PFI, and the effect of the number of targets simultaneously invisible. We compared the amount of PFI per trial, duration per PFI, and total duration of PFI during either 0, 1, 2, 3, or 4 target PFI periods (**Supplementary Figure 5**, blue bars). We found that while zero target PFI periods were most frequent (occurring 6 times per trial), when targets did disappear, simultaneous 4 target disappearances were the most common (4 times per trial, compared to <2 times per trial for 1, 2, and 3 target disappearances). This interesting trend continued for both the duration per PFI (7 s per 0 target, <2 s for 1-3 targets, 5.5 s for 4 targets) and total duration of PFI (32 s for 0, <2 for 1-3, and 20 s for 4, respectively), showing that 4 target disappearances were the most common, disappeared for a longer duration, and greatest total duration per trial.

To investigate whether these trends obtained over four-target locations were likely to occur by chance, we performed a shuffling analysis (see methods, and Supplementary Figure 5) and recalculated PFI characteristics (PFI per trial, PFI duration, and total duration), as a function of the number of targets filled-in (nPFI).

**Supplementary Figure 5** (c-h, grey bars) displays the results of this analysis, showing the mean across all 1000 shuffled sets of data. In contrast to observed data, the shuffled data showed PFI for 1, 2, and 3 targets being more common (6, 7, and 6 times per trial respectively), than for zero and four targets (each occurring less than 4 times per trial). For PFI duration, especially durations for 4 target PFI were shorter in the shuffled data. Strikingly, and in direct opposition to our observed data, the total duration of 0, 1, 2, and 3 target PFI per trial was roughly equivalent in shuffled data (each ~14 s duration), with 4 target PFI occurring for the least amount of time (<10 s).

To statistically evaluate the differences in these trends, we compared the coefficient for the quadratic fit of our observed data to all the coefficients in our null distribution for shuffled data. For all PFI measures, the observed coefficient was outside the 95 % of the null distribution (corresponding to *p* <.05). For the PFI per trial and total duration per trial, the positive slope for the effect of nPFI in our observed data is in direct opposition to the distribution of quadratic coefficients in our shuffled data. Taken together, we interpret this result as evidence of a synergistic spatial-interaction between multiple PFI targets, which may be mediated by attentional grouping ^18,52^.

### Jackknife latency estimate confirms changes in background SNR precede target SNR during endogenous PFI

To investigate the timing of changes to target and surround SNR, we superimposed the time-course for disappearances and reappearances, aligning at button press and release. We then compared the magnitude of these SNR time-courses using consecutive paired samples *t*-tests (two-tailed), at each time point. We identified that significant changes to the magnitude of surround-SNR (f2) preceded changes to target-SNR (f1), by approximately 200 ms (one time-step in our moving window spectrogram).

Previous investigations of PFI have frequency-tagged only a single target ^53^, or only the surrounding background (^18,52^, leaving the temporal order of these dynamics unclear. Here, we demonstrate that an increase in surround-SNR precedes an increase in target-SNR by approximately 200 ms (**Figure 2**). This demonstration is an important advance, because surround-specific changes precede target-specific responses, and the target-specific response is in the opposite direction (i.e., increasing, not decreasing, during PFI). This pattern challenges models of PFI that assert that adaptation of target-sensitive neurons in early retinotopic visual cortex precedes an increase in surround-activity, and the subsequent phenomenon of filling-in ^7,54,55^.

To confirm the differences between target and surround-specific changes in SNR, we also performed a non-parametric jackknife resampling procedure. As the waveforms under consideration for f1 and f2 have different peak amplitudes and shape, we avoided latency estimates based on peak-criterion (cf. Miller et al., 1998; 2009). Instead, we repeated our temporal cluster based analysis after subsampling participants in a leave-one-out analysis, to estimate the reliability of this effect. Specifically, we compared the SNR time-courses for disappearance and reappearance using running paired samples *t*-tests (two-tailed), at each time point. The first time-point in a temporally contiguous cluster of at least 500 ms was retained as the first difference in SNR during PFI. To increase the temporal resolution of our jackknife estimates, we linearly interpolated the SNR time-courses from 250 ms time-steps (4 Hz), to 1 ms time steps (1000 Hz)

The median, jackknifed estimates for these first significant time-points replicated the pattern observed in our group-level data (**Supplementary Figure 6**). The median first significant time-point for f2 (−1.22 s, SD = −0.02) preceded f1 (−0.43 s; SD = 0.26 s). Importantly, for every jackknifed subsample, changes in f2 always preceded f1, and the difference between these distributions was significant (Wilcoxon signed rank test; z= 3.52, *p* = 4.38 x10^−4^).

---

**Supplementary Figure 1.**
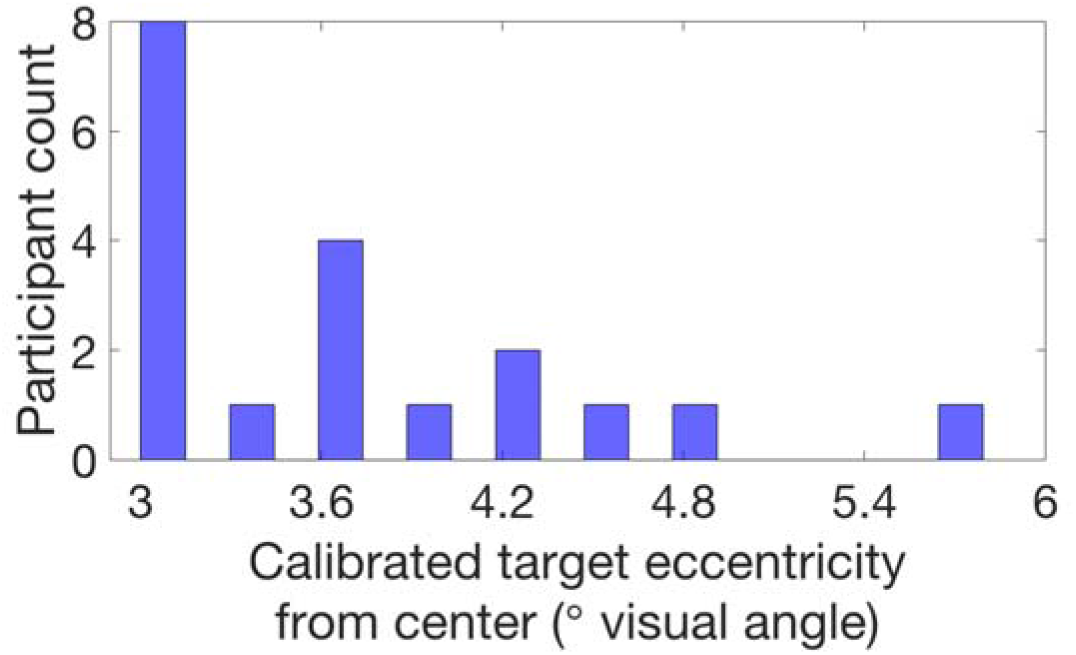
Calibrated target eccentricity. Across all participants, the final target eccentricity used throughout each experiment is shown.

**Supplementary Figure 2.**
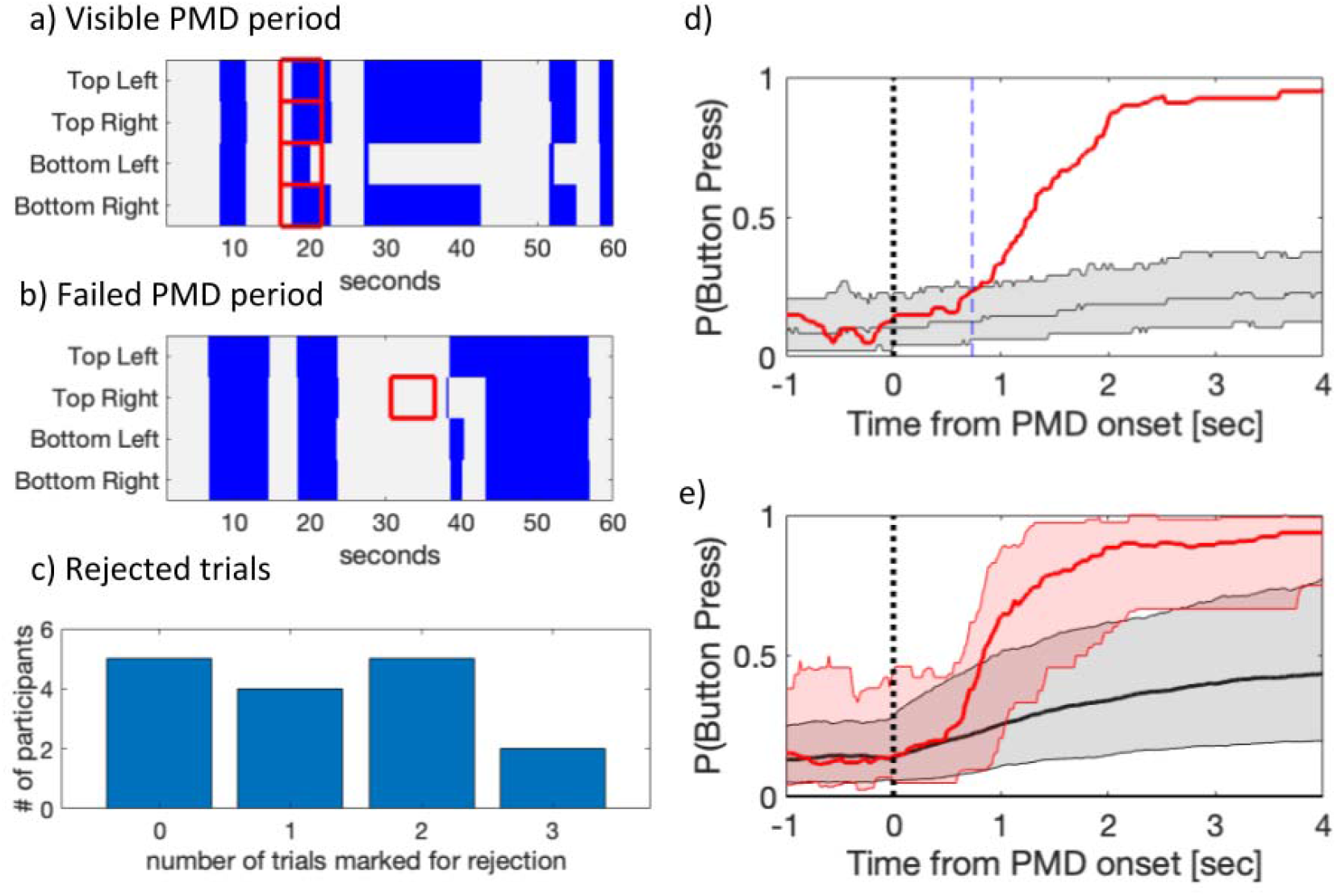
Behavioural responses during phenomenally matched disappearance (PMD). a-b) Example button-press responses during a 1-minute trial, blue area indicates button press periods (target disappearance) at each target location. PMD periods (indicated by red squares) were correctly detected in a) but not in b). c) A histogram count for the number of trials rejected across participants, when PMD onsets were not correctly detected. d) The time course of the button press likelihood around PMD-onset, and its associated null distribution for a single participant. The red line is the median time course for all PMD-onset across 48 trials. The black line and shading indicates the median and the 95% CI for the distribution across 1000 bootstraps, respectively (see Methods). We define the first time point that the observed button press time course (red) exceeded the bootstrapped CI (grey) as the reaction time to report PMD (0.73 s for this participant, marked with a vertical blue dashed line). e) The population time course of the button press around PMD-onset and its associated null distribution. The red line is the median, across the median time courses from each of our N=16 retained participants. The thick black line and shading is also the median of the median, and the 5% and 95% CI estimate obtained after bootstrap within each participant (computed with logit transform and presented after reverse transform).

**Supplementary Figure 3.**
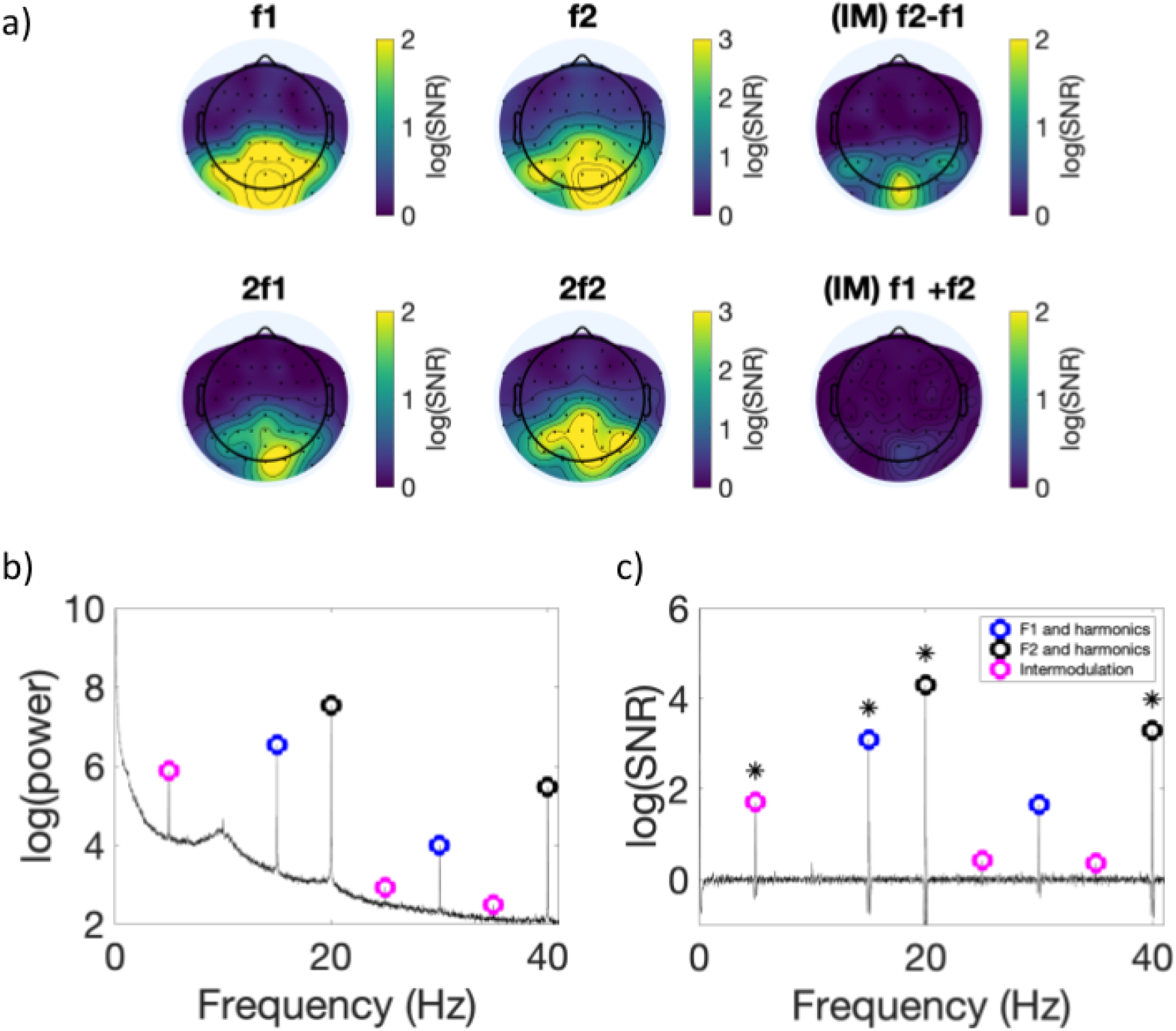
Whole trial SSVEP results. a) The topographic distribution of SNR across participants is shown at target (F1; 15 Hz), surround (F2; 20 Hz), intermodulation (F2-F1; 5 Hz), and higher harmonic (2f1; 2f2; f1+f2) frequencies. b) Average SSVEP responses and c) log (SNR) for all experimental periods in our paradigm (60 s whole-trials at channel POz). Target flicker and harmonics are shown in blue, surround flicker and harmonics are shown in black, with intermodulation components shown in magenta. In c), asterisks mark log(SNR) significantly different from 0 in spectrum at POz, FDR-adjusted across all frequencies to p < .05.

**Supplementary Figure 4.**
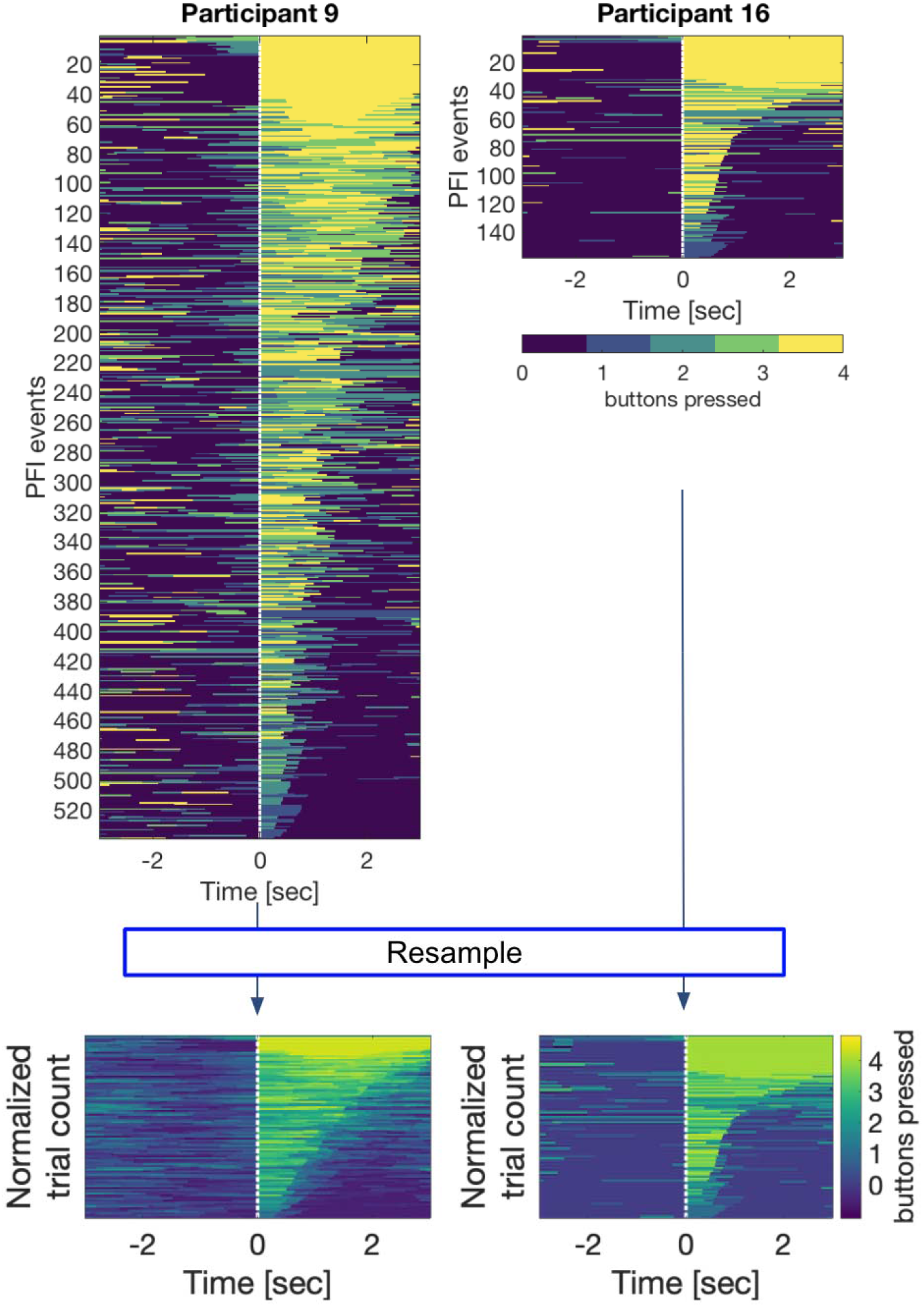
Preprocessing for event-by-event based image analyses. PFI events were first sorted according to “the amount of PFI” (the integral of the number of buttons pressed over 3 s) occurring after button-press or before button release. Each image along the y-axis was then resampled to normalize the number of trials from 1 to 100 samples. The same process was also applied to RESS log(SNR) after sorting by the amount of PFI per trial based on button press (or release). This image-based analysis enables us to compare PFI dynamics despite differences in the number of PFI events per participant.

**Supplementary Figure 5.**
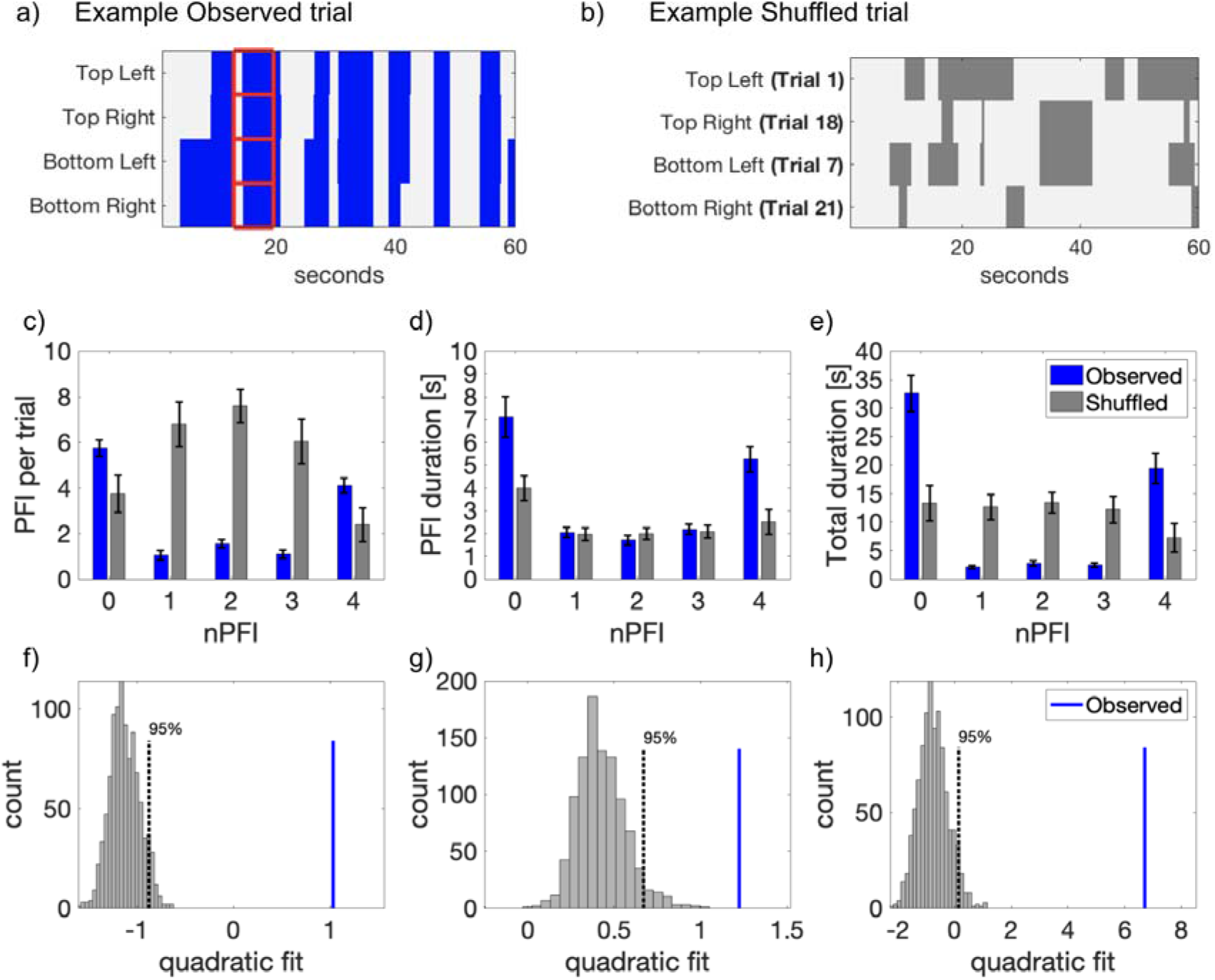
Behavioural data comparing PFI characteristics based on the number of targets perceptually filled in (nPFI) a) Example PFI data from one participant, displaying synergistic PFI across multiple locations. b) Example of shuffled data to test whether synergistic PFI occurs by chance. c) Instances of PFI per trial, d) mean duration per PFI, and e) total duration of PFI as a function of nPFI. All panels display both observed (blue) and shuffled (grey) data. For the observed data, error bars represent 1 SEM, corrected for within-participant comparisons ^40^. For the shuffled data, we first computed the SEM for each of the 1000 shuffled data sets. Then, as the error bar, we display the mean of these 1000 SEMs. f-h) 2nd order polynomial coefficients, for a quadratic fit for each of the PFI variables in c-e as a function of nPFI for observed (blue line) vs shuffled data (1000 sets, grey histogram).

**Supplementary Figure 6.**
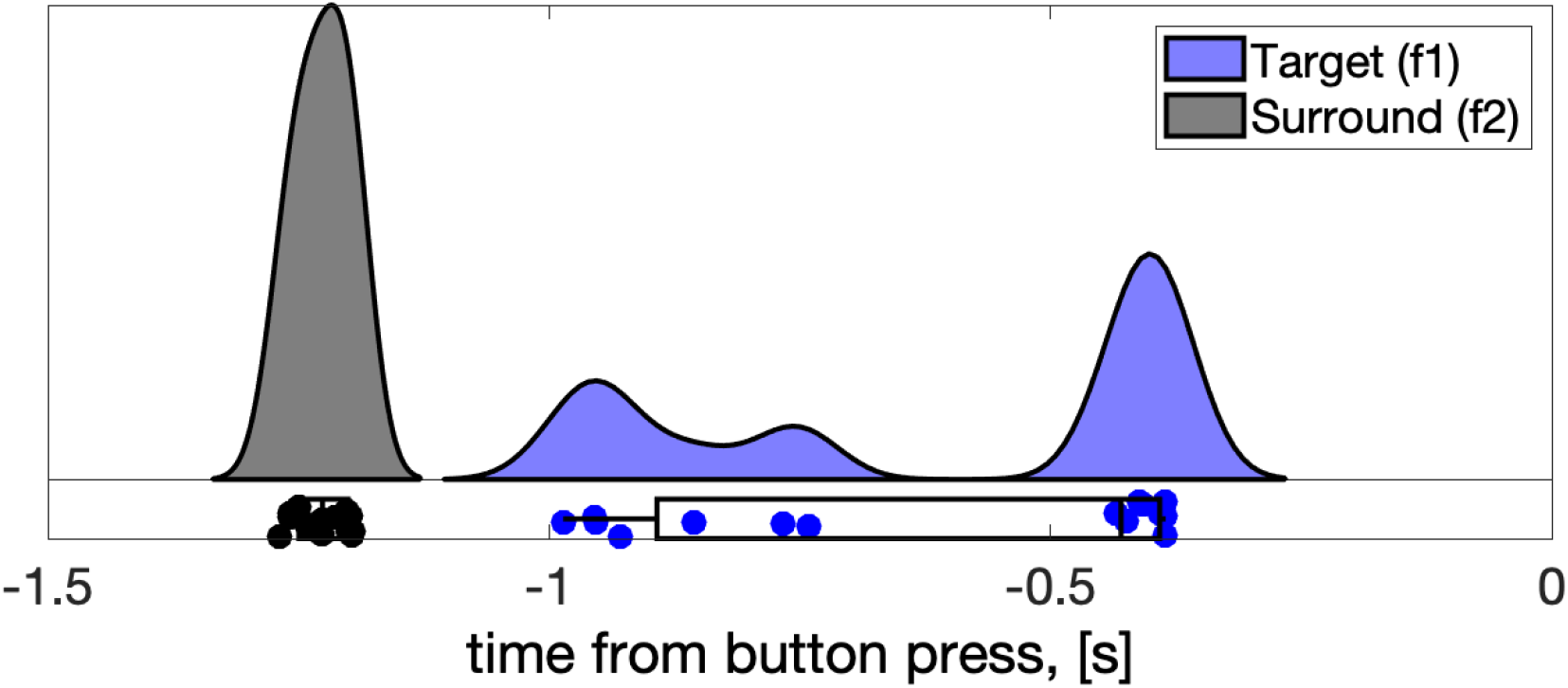
Jackknife analysis. First significant time-points in surround-specific SNR (f2; black), and target-specific SNR (f1; blue), after subsampling participants in a leave-one-out analysis.

## Notes

https://osf.io/hs7fn/

### Competing Interest Statement

The authors have declared no competing interest.

### Summary of Updates

Updated formatting

## References

1. Brown, R. J. & Norcia, A. M. A method for investigating binocular rivalry in real-time with the steady-state VEP. Vision Res. 37, 2401–2408 (1997).

2. Lansing, R. W. Electroencephalographic correlates of binocular rivalry in man. Science 146, 1325–1327 (1964).

3. Cobb, W. A., Morton, H. B. & Ettlinger, G. Cerebral potentials evoked by pattern reversal and their suppression in visual rivalry. Nature 216, 1123–1125 (1967).

4. Lou, L. Selective peripheral fading: evidence for inhibitory sensory effect of attention. Perception 28, 519–526 (1999).

5. De Weerd, P., Smith, E. & Greenberg, P. Effects of selective attention on perceptual filling-in. J. Cogn. Neurosci. 18, 335–347 (2006).

6. De Weerd, P., Desimone, R. & Ungerleider, L. G. Perceptual filling-in: a parametric study. Vision Res. 38, 2721–2734 (1998).

7. De Weerd, P., Gattass, R., Desimone, R. & Ungerleider, L. G. Responses of cells in monkey visual cortex during perceptual filling-in of an artificial scotoma. Nature 377, 731–734 (1995).

8. Dennett, D. C. Consciousness Explained. Boston (Little, Brown and Co) 1991. (1991).

9. Kingdom, F. & Moulden, B. Border effects on brightness: A review of findings, models and issues. Spat. Vis. 3, 225–262 (1988).

10. Weil, R. S. & Rees, G. A new taxonomy for perceptual filling-in. Brain Res. Rev. 67, 40–55 (2011).

11. Davidson, M. J., Graafsma, I. L., Tsuchiya, N. & van Boxtel, J. A multiple-response frequency-tagging paradigm measures graded changes in consciousness during perceptual filling-in. Neurosci Conscious 2020, (2020).

12. Vialatte, F.-B., Maurice, M., Dauwels, J. & Cichocki, A. Steady-state visually evoked potentials: focus on essential paradigms and future perspectives. Prog. Neurobiol. 90, 418–438 (2010).

13. Norcia, A. M., Appelbaum, L. G., Ales, J. M., Cottereau, B. R. & Rossion, B. The steady-state visual evoked potential in vision research: A review. J. Vis. 15, 4 (2015).

14. Morgan, S. T., Hansen, J. C. & Hillyard, S. A. Selective attention to stimulus location modulates the steady-state visual evoked potential. Proc. Natl. Acad. Sci. U. S. A. 93, 4770–4774 (1996).

15. Müller, M. M. et al. Feature-selective attention enhances color signals in early visual areas of the human brain. Proceedings of the National Academy of Sciences 103, 14250–14254 (2006).

16. Zhang, P., Jamison, K., Engel, S., He, B. & He, S. Binocular rivalry requires visual attention. Neuron 71, 362–369 (2011).

17. Müller, M. M. et al. Effects of spatial selective attention on the steady-state visual evoked potential in the 20-28 Hz range. Brain Res. Cogn. Brain Res. 6, 249–261 (1998).

18. Alais, D., Blake, R. & Lee, S.-H. Visual features that vary together over time group together over space. Nat. Neurosci. 1, 160–164 (1998).

19. Worden, M. S., Foxe, J. J., Wang, N. & Simpson, G. V. Anticipatory Biasing of Visuospatial Attention Indexed by Retinotopically Specific α-Bank Electroencephalography Increases over Occipital Cortex. J. Neurosci. 20, RC63–RC63 (2000).

20. Sokoliuk, R. et al. Two Spatially Distinct Posterior Alpha Sources Fulfill Different Functional Roles in Attention. J. Neurosci. 39, 7183–7194 (2019).

21. Foxe, J. J. & Snyder, A. C. The Role of Alpha-Band Brain Oscillations as a Sensory Suppression Mechanism during Selective Attention. Front. Psychol. 2, 1–13 (2011).

22. Foxe, J. J., Simpson, G. V. & Ahlfors, S. P. Parieto-occipital 1 0Hz activity reflects anticipatory state of visual attention mechanisms. Neuroreport 9, 3929–3933 (1998).

23. Gould, I. C., Rushworth, M. F. & Nobre, A. C. Indexing the graded allocation of visuospatial attention using anticipatory alpha oscillations. J. Neurophysiol. 105, 1318–1326 (2011).

24. Halgren, M. et al. The generation and propagation of the human alpha rhythm. Proc. Natl. Acad. Sci. U. S. A. 116, 23772–23782 (2019).

25. Jensen, O. & Mazaheri, A. Shaping functional architecture by oscillatory alpha activity: gating by inhibition. Front. Hum. Neurosci. 4, 186 (2010).

26. Macdonald, J. S. P., Mathan, S. & Yeung, N. Trial-by-Trial Variations in Subjective Attentional State are Reflected in Ongoing Prestimulus EEG Alpha Oscillations. Front. Psychol. 2, 1–16 (2011).

27. Maris, E. & Oostenveld, R. Nonparametric statistical testing of EEG- and MEG-data. J. Neurosci. Methods 164, 177–190 (2007).

28. Sassenhagen, J. & Draschkow, D. Cluster-based permutation tests of MEG/EEG data do not establish significance of effect latency or location. Psychophysiology 56, e13335 (2019).

29. Mehta, N. & Mashour, G. A. General and specific consciousness: a first-order representationalist approach. Front. Psychol. 4, 407 (2013).

30. Zeki, S. The disunity of consciousness. in Progress in Brain Research (eds. Banerjee, R. & Chakrabarti, B. K.) vol. 168 11–268 (Elsevier, 2007).

31. Mashour, G. A., Roelfsema, P., Changeux, J.-P. & Dehaene, S. Conscious Processing and the Global Neuronal Workspace Hypothesis. Neuron 105, 776–798 (2020).

32. van Vugt, B. et al. The threshold for conscious report: Signal loss and response bias in visual and frontal cortex. Science 360, 537–542 (2018).

33. Tononi, G. An information integration theory of consciousness. BMC Neurosci. 5, 42 (2004).

34. Oizumi, M., Albantakis, L. & Tononi, G. From the phenomenology to the mechanisms of consciousness: Integrated Information Theory 3.0. PLoS Comput. Biol. 10, e1003588 (2014).

35. Tononi, G., Boly, M., Massimini, M. & Koch, C. Integrated Information Theory: from consciousness to its physical substrates. Nat. Rev. Neurosci. in press, 450–461 (2016).

36. Haun, A. M. et al. Conscious Perception as Integrated Information Patterns in Human Electrocorticography. eNeuro 4, (2017).

37. van Boxtel, J. J. A., Tsuchiya, N. & Koch, C. Consciousness and attention: on sufficiency and necessity. Front. Psychol. 1, 1–13 (2010).

38. Tsuchiya, N. & Koch, C. The Relationship Between Consciousness and Top-Down Attention. in The Neurology of Conciousness (Second Edition) (eds. Laureys, S., Gosseries, O. & Tononi, G.) 71–91 (Academic Press, 2016).

39. Merten, K. & Nieder, A. Active encoding of decisions about stimulus absence in primate prefrontal cortex neurons. Proc. Natl. Acad. Sci. U. S. A. 109, 6289–6294 (2012).

40. Cousineau, D. Confidence intervals in within-subject designs: A simpler solution to Loftus and Masson’s method. Tutor. Quant. Methods Psychol. 1, 42–45 (2005).

41. Delorme, A. & Makeig, S. EEGLAB: an open source toolbox for analysis of single-trial EEG dynamics including independent component analysis. J. Neurosci. Methods 134, 9–21 (2004).

42. Bokil, H., Andrews, P., Kulkarni, J. E., Mehta, S. & Mitra, P. P. Chronux: a platform for analyzing neural signals. J. Neurosci. Methods 192, 146–151 (2010).

43. Oostenveld, R., Fries, P., Maris, E. & Schoffelen, J.-M. FieldTrip: Open source software for advanced analysis of MEG, EEG, and invasive electrophysiological data. Comput. Intell. Neurosci. 2011, 156869 (2011).

44. Kothe, C. A. & Makeig, S. BCILAB: a platform for brain-computer interface development. J. Neural Eng. 10, 056014 (2013).

45. Anstis, S. Adaptation to peripheral flicker. Vision Res. 36, 3479–3485 (1996).

46. Schieting, S. & Spillmann, L. Flicker adaptation in the peripheral retina. Vision Res. 27, 277–284 (1987).

47. Bigdely-Shamlo, N., Mullen, T., Kothe, C., Su, K.-M. & Robbins, K. A. The PREP pipeline: standardized preprocessing for large-scale EEG analysis. Front. Neuroinform. 9, 1–20 (2015).

48. Ding, J., Sperling, G. & Srinivasan, R. Attentional modulation of SSVEP power depends on the network tagged by the flicker frequency. Cereb. Cortex 16, 1016–1029 (2006).

49. Cohen, M. X. & Gulbinaite, R. Rhythmic entrainment source separation: Optimizing analyses of neural responses to rhythmic sensory stimulation. Neuroimage 070862 (2016).

50. Fujiwara, M. et al. Optokinetic nystagmus reflects perceptual directions in the onset binocular rivalry in Parkinson’s disease. PLoS One 12, e0173707 (2017).

51. Allen, M., Poggiali, D., Whitaker, K., Marshall, T. R. & Kievit, R. A. Raincloud plots: a multi-platform tool for robust data visualization. Wellcome Open Res 4, 63 (2019).

52. Davidson, M. J., Graafsma, I., Tsuchiya, N. & van Boxtel, J. J. A. Frequency-tagging visual background information enables multi-target perceptual filling-in to be distinguished from phenomenally matched replay. BioRxiv 499517 (2018).

53. Weil, R. S., Kilner, J. M., Haynes, J. D. & Rees, G. Neural correlates of perceptual filling-in of an artificial scotoma in humans. Proc. Natl. Acad. Sci. U. S. A. 104, 5211–5216 (2007).

54. Pessoa, L., Thompson, E. & Noë, A. Finding out about filling-in: a guide to perceptual completion for visual science and the philosophy of perception. Behav. Brain Sci. 21, 723–48; discussion 748–802 (1998).

55. Spillmann, L. & De Weerd, P. Mechanisms of surface completion: Perceptual filling-in of texture. Filling-in: From perceptual completion 81–105 (2003).

